# Optimal Field Sampling Framework for Disease Surveillance of Clonally Propagated Plants

**DOI:** 10.64898/2025.12.09.693344

**Authors:** Merveille Koissi Savi, Timothy A. Anake, Justin S. Pita

**Affiliations:** Université Félix Houphouët-Boigny, Central and West Africa Virus Epidemiology (WAVE), Bingerville, Côte d’Ivoire; Department of Mathematics, Covenant University, Canaan Land, Ota, Ogun State, Nigeria; Laboratoire d’Innovation pour la Santé des Plantes, UFR Biosciences, Université Félix Houphouët-Boigny

**Keywords:** Clonally propagated species, Delay Differential Equations, Surveillance program, Sample size estimation, Decision theory

## Abstract

Vegetatively propagated crops are vulnerable to pathogen accumulation, yet, surveillance strategies often fail to capture the interplay between local infection dynamics, spatial dispersal, and propagation practices. We develop a spatially explicit delay differential compartmental model that integrates within-farm transmission, between-farm dispersal, and cutting-based propagation. Analytical results demonstrate that the number of infected individuals increases sharply after the first replanting cycle due to latent infections in planting material. Embedding the model into a decision-theoretic sampling framework reveals a nonlinear relationship between prevalence and sampling effort. At epidemic onset, the required sampling intensity is higher than that at moderate to high prevalence levels. Simulation experiments demonstrate that adaptive sequential designs can reduce survey costs by up to 40% while maintaining detection probability, and spatial interpolation of prevalence improves outbreak delineation compared to uniform sampling. These findings highlight that effective early detection in clonal crops requires intensified sampling during low-prevalence phases, complemented by spatially targeted surveys. The coupled dynamical-statistical framework thus provides theoretical insights and operational tools for optimizing surveillance, enhancing outbreak detection, and guiding resource allocation in plant health programs.

**MSC codes:** 92D30, 92C60, 62P10

## 1. Introduction

Pests and diseases are among the most pressing threats to ongoing global efforts to achieve sustainable agriculture and address zero hunger world-wide. Specifically, losses incurred as a direct result of the impact of pests and diseases on crops, including tubers, roots, and bananas, have been estimated to be, on average, 17.2% for potatoes, 24% for cassava, and 29% for bananas [19, 16, 18, 3]. Tubers, roots, and bananas are integral to food security and are propagated vegetatively. They are particularly susceptible to disease because pathogens can persist and spread throughout the growing seasons. This is because farmers often source seeds from seemingly healthy plants. In many developing countries, seed exchange remains the primary means of acquiring planting material from one season to the next [1].

Effectively tackling these threats requires surveillance systems; yet, current approaches in plant health face challenges of ad hoc sampling strategies, limited resources, and difficulty balancing statistical power with costs and logistics. These challenges hinder adequate sampling at different spatial and temporal scales at the onset of, and in monitoring, the progression of the epidemic. Recent studies offer insights into optimizing surveillance strategies to address these issues. When resources are limited, a surveillance program under budget constraints is suggested, determining the intensity and frequency of inspections and sampling [13]. The authors developed a sequential adaptive delimiting survey with increasing spatial resolution for *Xylella fastidiosa* in Alicante, Spain. Optimization was carried out through simulation, and sampling intensity thresholds were evaluated based on their effects on incidence estimation. Although this method offers a structure for estimating samples, its precision is insufficient to reflect the vertical and horizontal transmission dynamics, as in mechanistic models. This case study demonstrates the majority of several sample size estimation methods, which are often not rooted in epidemiological theory, resulting in difficulties with adaptability and making decisions based on information. The integration of epidemiological perspectives has been shown to improve sampling initiatives, thereby facilitating a more efficient identification of emerging pathogens [17].

Mathematical models elucidate disease mechanisms in vegetatively propagated crops and evaluate the effectiveness of disease management strategies. Deterministic compartmental models, including susceptible, infected, and removed, are employed to analyze the transmission dynamics of cassava mosaic disease, which spreads through vectors and contaminated planting materials in cassava farms [5]. Specifically, the authors determined conditions for local (global) asymptotic stability and demonstrated that Hopf Bifurcation may occur when roguing is used to control the epidemics.

The unpredictable patterns of crop outbreaks associated with regional environmental factors was captured through isotropic kernel dispersal and stochastic processes, in which the transmission probabilities decay along all radial directions as one moves farther from the epidemic front [6]. This framework was implemented to assess different cassava brown streak disease (CBSD) management options and optimize disease surveillance protocols by examining the effects of survey timing, sampling intensity, and diagnostic accuracy on CBSD detection and prevalence estimation [7]. They found that a minimum of 30 plants need to be surveyed within a farm, approximately 299 days post-planting, to obtain a minimum accuracy of 50% precision in identifying CBSD. However, the accuracy of the CBSD prevalence within a farm remains challenging since its estimation depends primarily on survey accuracy rather than the number of plants surveyed or the timing of the survey. Moreover, for the successful early identification of CBSD, assessments must be conducted in areas lacking previous disease documentation, and the number of farms per region (per-region sample size) should align with the density of cassava farms [8]. Even though these previous studies explored and proposed a sampling estimation on the onset of an epidemic, their application is still constrained by the thorough definition of the host landscape. They fall short of establishing a universal framework applicable to other vegetatively propagated crops at the onset and monitoring of epidemics.

Moreover, many crop disease surveillance programs, especially in developing countries, operate under budget constraints, limiting the intensity and frequency of sampling. This necessitates sampling optimization strategies to maximize efficiency within these limitations while balancing the trade-off between statistical power, costs, and logistical constraints [2, 13]. Although optimizing sampling offers promising solutions, their implementation must consider local contexts and pathogen-specific dynamics.

We introduce a theoretical framework for sampling estimation. Specifically, we develop a dynamic model of disease spread in clonally propagated plants that captures delay and feedback mechanisms and local spatial variations in transmission patterns. We then derive a prevalence-based utility function that links epidemic dynamics to surveillance performance. Finally, we establish an optimization framework to determine sample sizes and strategies that achieve reliable detection within resource constraints during the onset and monitoring of an epidemic.

## 2. Methodology: Dynamical Model and Sampling Framework

This section presents a formulation of a spatially explicit, delay differential compartmental model capturing the transmission dynamics of a clonal plant pathogen across multiple farms. We integrate local transmission, spatial dispersal, and propagation through cuttings into a unified model.

### 2.1. Model Structure and Assumptions

The propagation of epidemics is based on the assumption of a vertical spread of the epidemics. The model incorporates delays and feedback. Thus, the seeds utilized at any given time are derived from the preceding period. Over time, the incidence of disease can increase. This indicates that in a plant population that is infected at a given time, the presence of asymptomatic plants can result in the subsequent generation of symptomatic plants. Conversely, we hypothesize that the population of each farm remains constant due to the absence of immigration or emigration of plants. Intra-farm transmission is modeled on a mass-action model, assuming random mixing and a large population. By contrast, between-farm transmission depends on the distance and density of the infected plant. The propagation through cuttings occurs regularly, with a fraction *γ* of plants replaced. The probability of transmitting infection through cuttings depends on the proportion 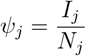 of symptomatic plants on the donor farm.

A fixed time delay *τ* is incorporated to represent the latent period between infection and the onset of symptoms, during which plants are not yet infectious.

We consider a meta-population system of *M* farms indexed by *j* = 1, …, *M*. Each farm consists of a closed plant population partitioned into three epidemiological compartments:

- *S*_*j*_(*t*): Number of *susceptible* plants;
- *L*_*j*_(*t*): Number of *latent* infected plants (infected but not yet infectious);
- *I*_*j*_(*t*): Number of *infectious* (symptomatic) plants;
- *N*_*j*_(*t*) = *S*_*j*_(*t*) + *L*_*j*_(*t*) + *I*_*j*_(*t*): Total plant population, assumed constant.

#### 2.1.1. Within-farm or Local Spread

The virus transmission within a farm occurs according to a mass-action model, where the infection rate in farm *j* depends on the density of infected plants within the farm. The infected plants cause new infections at a rate proportional to the number of susceptible and infected plants.

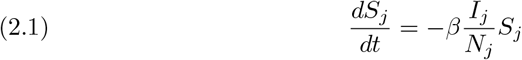

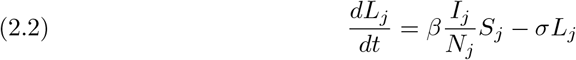

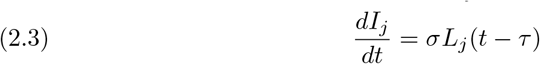

where *β* is the transmission rate within farm *j* and *σ* is the rate at which latent plants develop symptoms and become infectious; *τ* is a fixed delay representing the latent period before plants become infectious.

#### 2.1.2. Cutting-Based Spread

The infected plants can be used for propagation, but only symptomatic (*I*) plants contribute to transmission via cuttings. If the farmer randomly selects cuttings, the probability of choosing an infected cutting is proportional to the fraction of symptomatic plants.

Let *γ* be the fraction of new plants generated per time step. Besides, let us assume that the cutting behavior changes depending on local infection awareness. We define a feedback function *f* (*ψ*_*j*_) that reduces the use of infected cuttings as awareness increases. Thus

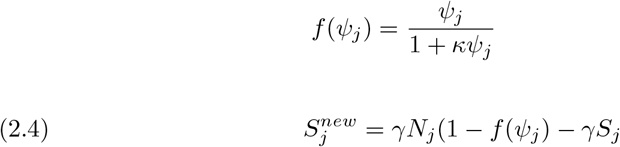

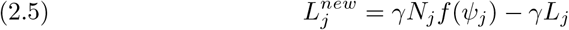

where 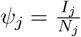 The fraction *ψ*_*j*_ of new plants comes from infected mothers and starts in the latent stage (*L*_*j*_). The remaining fraction (1 − *ψ*_*j*_) is healthy and joins *S*.

#### 2.1.3. Between-Farm Spread (spatial dynamics)

The infection also spreads between farms. We assume that the infection pressure from farm *j* to farm *k* depends on the distance between farms *d*_*jk*_ and the number of infected plants at farm *j*. The spatial spread follows a distance-decay model, meaning the probability of infection decreases with increasing distance between farms. The rate of spread from farm *j* to farm *k* is:

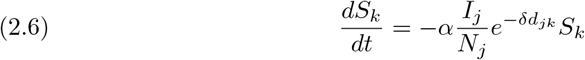

where *α* is the scaling parameter for between-farm infection, δ is the distance decay constant, *d*_*jk*_ is the geographic distance between farms *j* and *k*

The complete meta-population system can be obtained by combining the local and spatial spread.

#### 2.1.4. Complete System with Cutting-Based Spread

Combining within-farm infection, between-farm spread, and propagation through cuttings, the complete system for each farm *j* is:

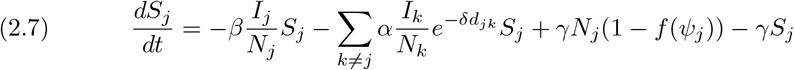

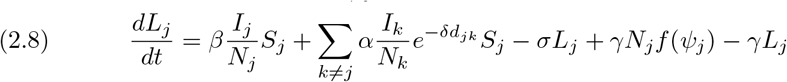

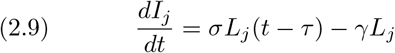

### 2.2. Mathematical Properties of the System

#### 2.2.1. Well-posedness and Positivity

##### Theorem 1

Let the initial history *x*_*j*_(*t*) = (*S*_*j*_(*t*), *L*_*j*_(*t*), *I*_*j*_(*t*)) be continuous and non-negative for *t* ∈ [−*τ*, 0], and all parameters non-negative. Then the system admits a unique, non-negative, and bounded solution for all *t* ≥ 0.

*Proof*: Existence and uniqueness follow from the standard theory of functional differential equations [9] under Lipschitz continuity. Non-negativity is preserved since each compartmental equation includes non-negative inflow and outflow terms that vanish at zero. Summing the three equations shows *N*_*j*_ is conserved.

##### Invariant Region

We define the biologically feasible region Ω_*j*_ for each farm *j* as:

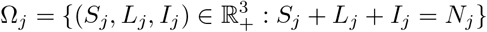

###### Theorem 2

If *x*_*j*_(0) ∈ Ω_*j*_ for all *j*, then *x*_*j*_(*t*) ∈ Ω_*j*_ for all *t* ≥ 0.

#### 2.2.2. Disease-free Equilibrium

##### Theorem 3

The disease-free equilibrium (DFE) occurs when *I*_*j*_ = 0, *L*_*j*_ = 0, and *S*_*j*_ = *N*_*j*_ for all *j*. The system is at rest, and no new infections occur in this state.

*Proof:* Set *I*_*j*_ = 0 and *L*_*j*_ = 0 for all *j*. Then 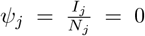, and *f*(*ψ*_*j*_) = *f*(0) = 0. Substituting into the system:

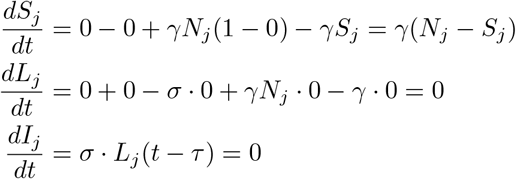

Thus, 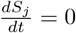 if and only if *S*_*j*_ = *N*_*j*_. Hence, the DFE is given by:

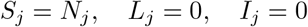

and the system remains in this state, confirming it is an equilibrium. To ensure equilibrium at the DFE, we must have:

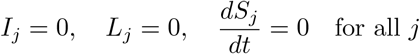

Substituting into the system:

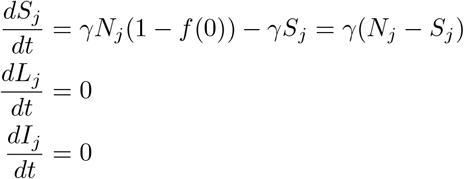

Thus, the DFE is given by:

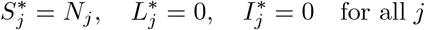

##### Remark

To ensure that the system is at equilibrium when *I*_*j*_ = 0, we assume that either the propagation term *γ* is turned off in the absence of infection (e.g., *γ* = 0 when *I*_*j*_ = 0), or we accept that the DFE corresponds to a state where all *S*_*j*_ = *N*_*j*_, that is, no new propagation or replacement occurs. Aligning with the assumption that cutting-based propagation is disease-driven.

#### 2.2.3. Stability Analysis of the DFE

To analyze the local stability of the DFE, we linearize the system around:

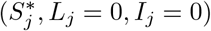

Define the state vector:

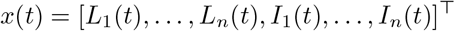

Let 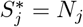 at the DFE. Then, the linearized equations for each patch *j* are:

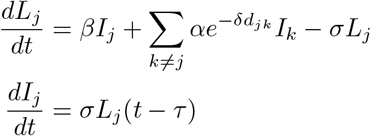

Hence, the linear system:

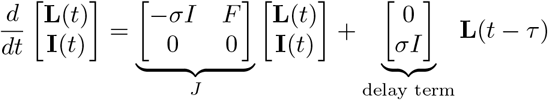

where *F* is the matrix encoding transmission from infectious individuals to latents:

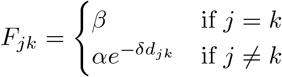

This delayed system’s stability can be analyzed via the next-generation matrix (NGM). Let *F* denote the infection matrix and *V* the transition matrix. Then the NGM is:

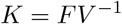

The spectral radius of *K*, denoted by *R*_0_, determines the stability of the DFE:

- If ℛ _0_ *<* 1, the DFE is locally asymptotically stable.
- If ℛ _0_ *>* 1, the DFE is unstable and infection can invade.

The DFE is locally asymptotically stable if the basic reproduction number ℛ _0_ *<* 1, meaning that on average, each infected plant causes less than one new infection across the system.

#### 2.2.4. Determination of the Basic Reproduction Number *R*_0_

The basic reproduction number *R*_0_ is computed using the *Next Generation Matrix (NGM)* vandendriessche2002. Let’s identify the infected compartments and the transmission structure of the model.

The system of equations is given by Equations (2.7), (2.8) and (2.9)

Next, the focus is on the infection subsystem involving the latent and infectious compartments *L*_*j*_ and *I*_*j*_.

##### Infection and Transition Terms

Let the vector of infected states be **x** = (*L*_*j*_, *I*_*j*_). We define the vector of new infections ℱ _*j*_ and the transitions 𝒱_*j*_ as follows:

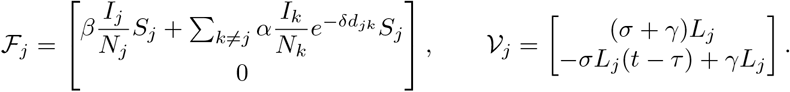

At the disease-free equilibrium (DFE), we assume *S*_*j*_ = *N*_*j*_, *L*_*j*_ = *I*_*j*_ = 0, and *f* (*ψ*_*j*_) govern only demographic inputs.

##### Jacobian Matrices

Let *F* = *D*_*x*_ℱ and *V* = *D*_*x*_𝒱 evaluated at the DFE. We are interested in the new infection terms into the *L*_*j*_ compartment. The entries of the Jacobian *F* at DFE are:

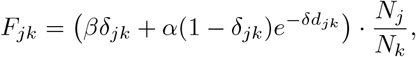

where δ_*jk*_ is the Kronecker delta.

The transition matrix *V* is diagonal with entries:

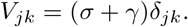

##### Next Generation Matrix and *R*_0_

The Next Generation matrix is given by:

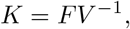

with components:

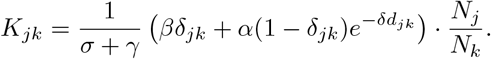

The basic reproduction number *R*_0_ is the spectral radius (i.e., the dominant eigen-value) of the matrix *K*:

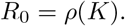

This expression captures both local and spatially mediated transmission, modulated by distance-dependent decay and normalized by the total population sizes at each site.

## 3. Sampling Framework and Surveillance Utility

We couple the metapopulation dynamics to a statistically explicit observation model and an adaptive design. Goals are: (i) maximize early outbreak detection, (ii) minimize uncertainty in prevalence/forecasts, and (iii) reallocate effort as new data arrive.

### 3.1. Observation Model

To avoid overloading notation, we reserve *ψ*_*j*_(*t*) = *I*_*j*_*/N*_*j*_ for the cutting-feedback in the process model. For sampling and surveillance, we instead work with *compartment fractions*, i.e., the proportions of plants in each epidemiological class at site *j*:

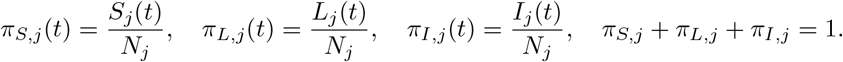

For a detection channel *c* (e.g. visual inspection or PCR testing), the probability of correctly identifying infection depends on the test’s sensitivity and specificity. We denote *Se*_*c,L*_ as the sensitivity for detecting latent plants, *Se*_*c,I*_ as the sensitivity for detecting infectious plants, and *Sp*_*c*_ as the specificity of the test (probability of correctly classifying a susceptible plant as negative). The effective probability that a plant drawn from site *j* at time *t* is classified as positive is:

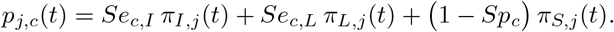

The number of observed positives, *Y*_*j,c*_(*t*), when *n*_*j,c*_(*t*) plants are sampled at site *j* using channel *c*, follows a binomial distribution:

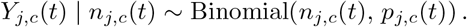

Visual inspection typically has *Se*_visual,*L*_ = 0 (unable to detect latent infections), while PCR usually achieves *Se*_PCR,*L*_ *>* 0 (able to detect latent).

### 3.2. Design Variables, Budget and Routing

The design problem involves choosing sampling intensities and locations. The decision variables are:

- *n*_*j,c*_(*t*): the number of samples taken at site *j*, time *t*, using channel *c*,
- *A*(*t*) ⊂ *{*1, …, *M}*: the set of sites visited at time *t*.

*λ*_*c*_ is the per-sample cost for channel *c*, and *C*(*A*(*t*)) is the routing or fieldwork cost for visiting the set of sites *A*(*t*). Given a total budget *B*, the cost constraint is:

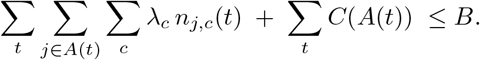

### 3.3. Detection Probability and Sample Size

To guarantee outbreak detection, we calculate the minimum number of samples *n*_*j,c*_(*t*) needed to detect at least one positive with probability *γ*. The required sample size is:

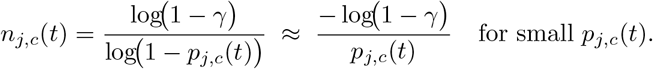

This formula automatically accounts for imperfect sensitivity and specificity through *p*_*j,c*_(*t*) and improves on heuristic formulas that use only *ϕ*_*j*_ = (*L*_*j*_ + *I*_*j*_)*/N*_*j*_.

### 3.4. Spatial Correlation and Effective Sample Size

Plants sampled within the same site may not be independent, leading to correlation among observations. Let *ρ*_*j,c*_ denote the within-site correlation for site *j* and channel *c*. The *effective sample size* is reduced to:

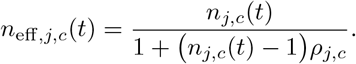

### 3.5. Utilities Grounded in Process and Observation

Two main objectives drive the definition of surveillance utilities: early detection and information for monitoring/inference.

#### Early detection (risk-weighted hit probability)

Here, the utility is the expected benefit of detecting infection at site *j*, weighted by the site-specific risk *w*_*j*_(*t*), minus the sampling cost:

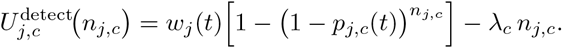

#### Monitoring and Inference (information-based)

Let *θ* denote epidemiological process parameters (e.g. *β, α, δ, σ, τ, γ, κ*) or a derived target such as prevalence Φ(*t*). The Fisher information measures how much precision about *θ* is gained from observing samples at site *j* and channel *c*:

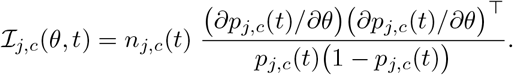

- Optimality criteria guide the design:
- *A*-optimality minimizes the trace of the covariance matrix of estimated parameters,
- *D*-optimality maximizes the determinant of the Fisher information matrix, forecast-targeted utility maximizes the expected reduction in uncertainty for a future prevalence Φ(*t* + Δ).

### 3.6. Optimization with Constraints

The overall design problem is to maximize total utility (across all times, sites, and channels) subject to budget constraints:

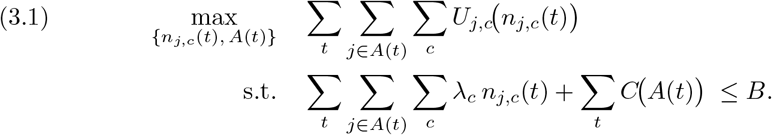

This formulation makes the *value of information* explicit: sampling designs are judged by their ability to improve detection probability or reduce uncertainty about key epidemiological parameters and forecasts, relative to their costs.

### 3.7. Adaptive Sequential Sampling

The adaptive sequential sampling framework is designed to jointly forecast epidemic dynamics and optimize surveillance strategies under budgetary and logistical constraints. At each decision point, the metapopulation model generates predictions of the compartmental probabilities (*π*_*S,j*_, *π*_*L,j*_, *π*_*I,j*_), providing the expected prevalence *p*_*j,c*_(*t*) for each location and crop. These forecasts are then used to identify the optimal allocation of resources *A*(*t*) and sampling intensities *{n*_*j,c*_(*t*)*}*, ensuring that inspection efforts remain cost-effective and operationally feasible. Data collected from the field, denoted *Y*_*j,c*_(*t*), are incorporated through the observation model to update epidemic states and model parameters, thereby refining predictions and allocation decisions. The process is repeated at each subsequent interval *t* + Δ, creating an iterative cycle that continuously adapts surveillance to the evolving epidemic.

### 3.8. Regional Prevalence and Targets

The *regional prevalence* Φ(*t*) is defined as the weighted average proportion of infected plants (latent or infectious) across all *M* sites:

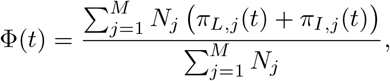

where *N*_*j*_ is the total number of plants at site *j, π*_*L,j*_(*t*) = *L*_*j*_(*t*)*/N*_*j*_ is the latent fraction, and *π*_*I,j*_(*t*) = *I*_*j*_(*t*)*/N*_*j*_ is the infectious fraction. This aggregate measure can be used directly as a monitoring or forecast target within the information-based utility framework.

## 4. Optimization Framework: Theoretical Computation

Let *θ* = (*β, α, δ, σ, τ, γ, κ*)^⊤^ denote the epidemiological parameter vector, where *β* is the transmission rate, *α* is the coupling rate from external sources, δ is the spatial decay parameter, *σ* is the progression rate from latent to infectious, *τ* is the latent period (delay), *γ* is the removal/replanting rate, and *κ* is the behavioral feedback strength. The metapopulation delay differential equation (DDE) model given in Eqs. (2.7)–(2.9) defines the compartment fractions *π*_*S,j*_(*t*), *π*_*L,j*_(*t*), *π*_*I,j*_(*t*) and the channel-specific detection probability.

For a detection channel *c* (e.g. visual inspection, PCR), with sensitivities *Se*_*c,L*_, *Se*_*c,I*_ and specificity *Sp*_*c*_, the probability of observing a positive test at site *j* and time *t* is

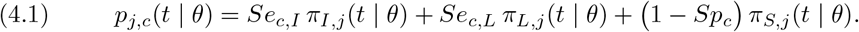

Given a per-site/time sample size *n*_*j,c*_(*t*) ∈ ℕ, the observation model is

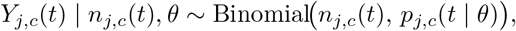

where *Y*_*j,c*_(*t*) is the number of positives observed.

### 4.1. Pre-Optimization: Global Sensitivity and Design Priors

#### Global screening

Let Θ ⊂ ℝ^7^ be a hyper-rectangle representing plausible parameter ranges, e.g.

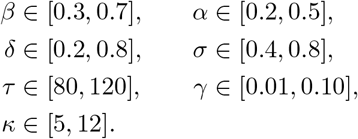

Draw an *N*-point Latin Hypercube sample (LHS) 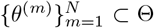 and compute scalar targets such as

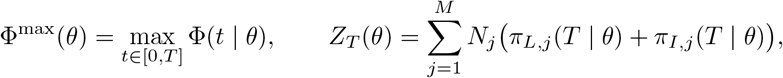

where Φ^max^(*θ*) is the maximum prevalence over the horizon and *Z*_*T*_ (*θ*) is the total number of infected plants at the final time *T*.

Define the partial rank correlation coefficient (PRCC) of a scalar target *g*(*θ*) with respect to parameter *θ*_*k*_:

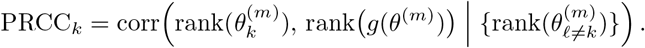

Parameters with |PRCC_*k*_| exceeding a threshold (e.g. 0.4) are deemed influential and retained with informative priors, while the remainder receive diffuse priors. This yields a *design prior π*(*θ*) supported on Θ.

#### Detection sample size lower bound

Fix site *j*, channel *c*, and time *t*. The probability of detecting at least one positive sample is

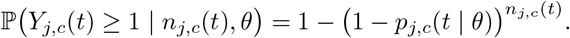

To guarantee a target detection probability *γ*^⋆^ ∈ (0, 1) *for all θ* in an uncertainty set Θ^∗^ ⊆ Θ, a sufficient condition is

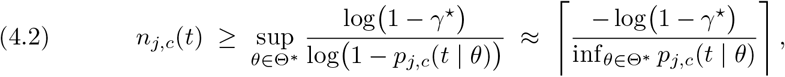

where the approximation uses log(1 − *x*) ≈ −*x* for small *x*.

### 4.2. During Optimization: Robust Utility Maximization

Let *A*(*t*) ⊂ *{*1, …, *M}* denote the set of visited sites at time *t, λ*_*c*_ *>* 0 the per-sample cost for channel *c*, and *C*(*A*(*t*)) the routing/field cost for visiting sites.

We define a *risk-weighted detection utility*:

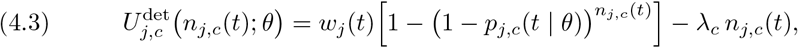

where *w*_*j*_(*t*) is a site-specific weight (e.g. economic or epidemiological importance).

An *information utility* is defined based on Fisher information for a scalar target parameter *ϑ* = *𝓁*^⊤^*θ*, with *𝓁* a projection vector:

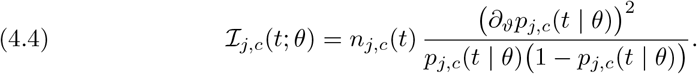

Let Π denote the design prior from the pre-optimization stage. The robust Bayes objective is then

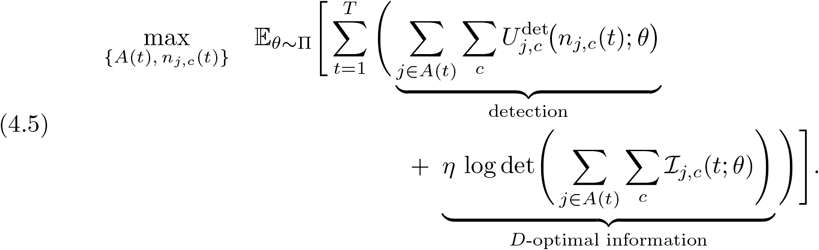

subject to the budget constraint

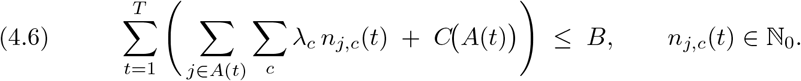

Here *η* ≥ 0 balances detection utility against information gain. The expectation is approximated by Monte Carlo integration over Π.

#### Marginal allocation condition (continuous relaxation)

For fixed *A*(*t*), *θ*, and treating *n*_*j,c*_(*t*) as continuous, the optimal *n*_*j,c*_(*t*) for the detection utility satisfies

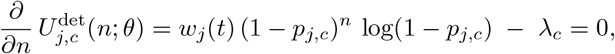

yielding

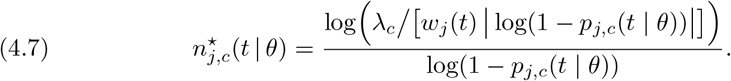

In the robust setting, replace *p*_*j,c*_(*t* | *θ*) with its effective value under Π, such as 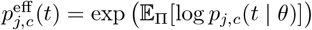 or the worst-case 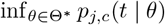.

#### Feasibility via lower bounds

A design is feasible if (4.6) holds and 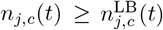, with 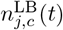 given by (4.2) for a program-specified target *γ*^⋆^ and uncertainty set Θ^∗^.

### 4.3. Post-Optimization: Local Sensitivity, Min–Max Robustness, and Expected Value of Perfect Information (EVPI)

#### Local sensitivity of the chosen design

Given an optimal design 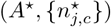, robustness is assessed by perturbing influential parameters *θ*_*k*_ one-at-a-time:

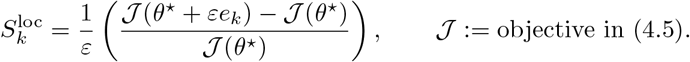

Here *e*_*k*_ is the unit vector in direction *k* and *ε* is a small perturbation size. Small 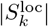 indicates robustness to *θ*_*k*_.

#### Worst-case (min–max) guarantee

Define the uncertainty set 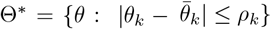 around a nominal 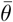. The min–max performance of the design is

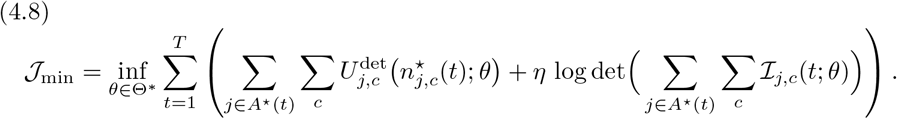

A design is *ξ-robust* if 𝒥_min_ ≥*ξ* 𝔼 _Π_[𝒥] for some *ξ* ∈ (0, 1) (typically *ξ* ≈ 0.8–0.9).

#### Value of information (EVPI/Expected Value of Sample Information (EVSI))

Let 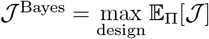 and 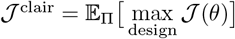. The expected value of perfect information is

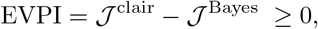

quantifying the benefit of resolving all uncertainty in *θ* before design. For a proposed pilot design *d*_pilot_, the expected value of sample information is

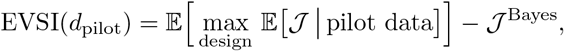

which ranks pilot allocations by their impact on the subsequent optimal design.

## 5. Spatially Explicit Sampling Optimization with Bayesian Sequential Updating

We explicitly model the sampling allocation problem over a rasterized spatial domain to incorporate spatial structure into surveillance. The study area is partitioned into square cells of resolution 5 km*×*5 km, consistent with operational sampling grids. Each cell *u* is assigned a prevalence weight *π*_*u*_(*t*) derived from the mechanistic epidemic model (Eqs. 2.7–2.9) and smoothed using inverse distance weighting (IDW). Specifically, the prevalence field is estimated as

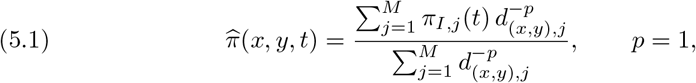

where *d*_(*x,y*),*j*_ denotes the Euclidean distance from spatial coordinate (*x, y*) to farm *j*. The resulting rasterized surface 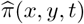 represents a smoothly interpolated prevalence map at 5 km resolution.

### Spatial coupling via dispersal kernel

Spatial dependence between neighboring cells is regulated by an isotropic discrete power-law dispersal kernel *K*(*d*) of the form

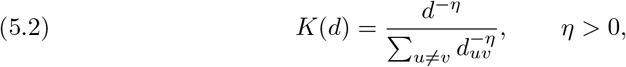

where *d*_*uv*_ is the distance between centroids of cells *u* and *v*, and *η* is the power-law exponent, which controls how steeply dispersal decays with distance. This kernel governs the probability of transmission between cells, linking prevalence in one location to the force of infection in its neighbors. In practice, the kernel ensures that high-prevalence foci exert spatial influence on surrounding areas with intensity decaying polynomially with distance.

### Bayesian sequential updating

For each cell *u*, let *π*_*u*_ denote the true but unknown prevalence. We place a Beta prior reflecting the IDW-smoothed prevalence estimate:

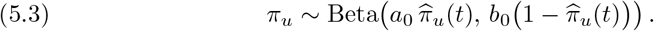

When *n*_*u*_ samples are collected and *y*_*u*_ positives observed, the posterior is

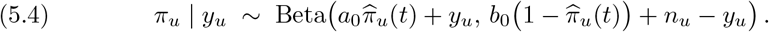

This posterior distribution dynamically updates prevalence estimates and propagates information spatially via the kernel *K*(*d*), ensuring that positive detections in one cell increase posterior prevalence in nearby cells.

### Sequential sampling rule

At each iteration *r*, the next sampling location *u*^⋆^ is chosen to maximize the expected surveillance utility:

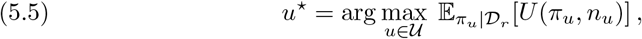

where 𝒟_*r*_ is the data collected up to round *r*, and *U* is a decision-theoretic utility function. Two regimes are emphasized:

1. **Early detection:** Utility is the probability of exceeding a prevalence threshold *π*_*u*_ *> π*_min_, targeting high-risk areas to maximize probability of outbreak detection.
2. **Epidemic monitoring:** Utility is the expected reduction in posterior variance, prioritizing allocation to regions where uncertainty about prevalence is highest.

### Integrated optimization

This Bayesian sequential framework links the mechanistic epidemic model, spatial interpolation via IDW, and dispersal coupling via the kernel *K*(*d*).

## 6. Parameterization and Simulation setup

Table 1 lists the baseline parameters used for numerical illustration together with literature sources or estimation notes. These values are consistent with clonal plant pathosystems (e.g., CBSD) and serve solely as a baseline for theoretical computations below.

**Table 1:**
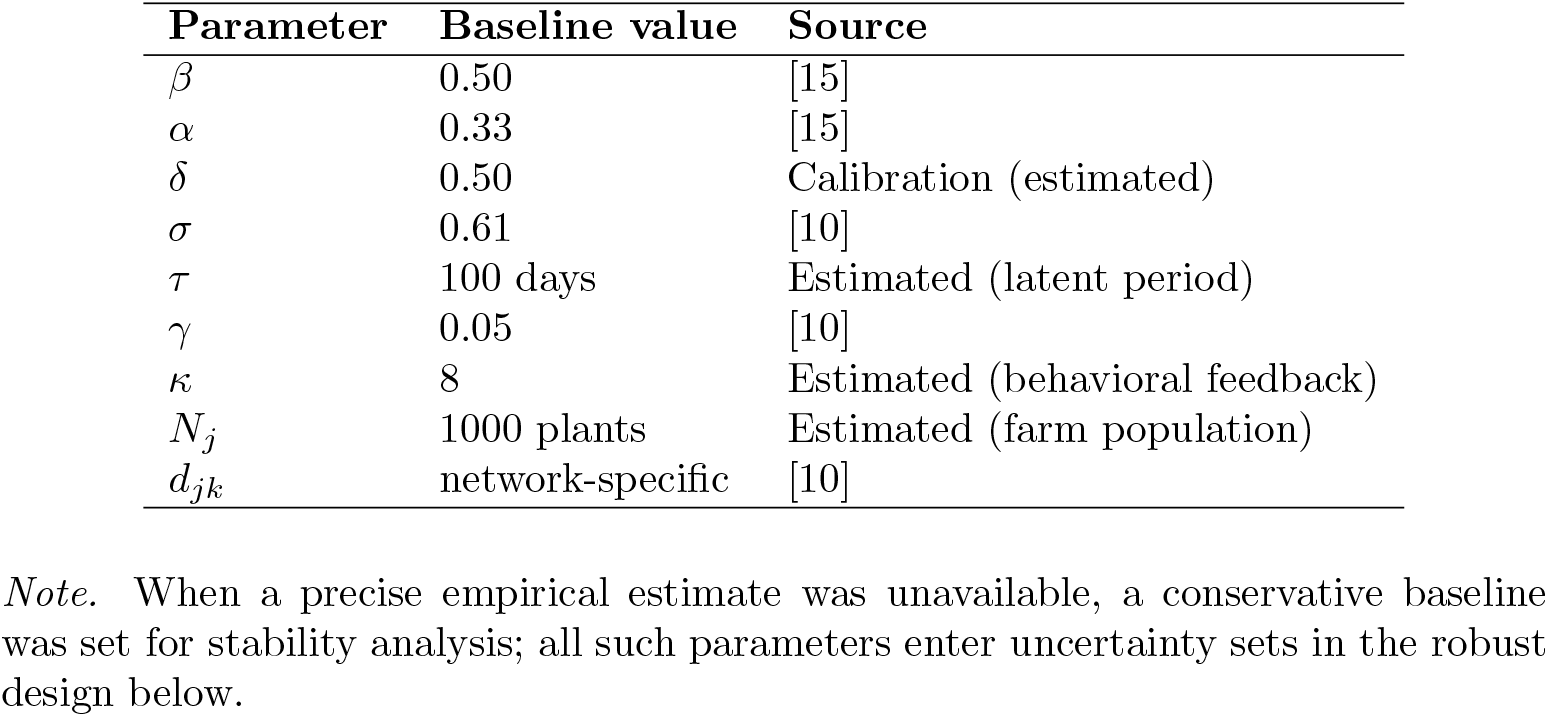
Baseline parameter values and sources used for theoretical computations.

| Parameter | Baseline value | Source |
| --- | --- | --- |
| $\beta$ | 0.50 | [15] |
| $\alpha$ | 0.33 | [15] |
| $\delta$ | 0.50 | Calibration (estimated) |
| $\sigma$ | 0.61 | [10] |
| $\tau$ | 100 days | Estimated (latent period) |
| $\gamma$ | 0.05 | [10] |
| $\kappa$ | 8 | Estimated (behavioral feedback) |
| $N_j$ | 1000 plants | Estimated (farm population) |
| $d_{jk}$ | network-specific | [10] |

*Note.* When a precise empirical estimate was unavailable, a conservative baseline was set for stability analysis; all such parameters enter uncertainty sets in the robust design below.

## 7. Numerical Resolution

The system of delay differential equations defined in Eqs. (2.7)–(2.9) does not admit a closed-form solution and must therefore be solved numerically. We adopted a fourth-order Runge-Kutta (RK4) scheme with a fixed time step Δ*t* = 0.1 days for the deterministic components, combined with a discrete-time delay approximation to account for the latent period *τ*. For state variables with delay terms, the method stores the history function over the interval [−*τ*, 0] and interpolates values at *t* − *τ* using linear interpolation.

Formally, for the vector of state variables

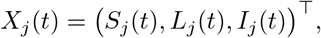

the system can be written as

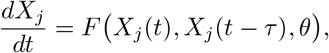

where *θ* = (*β, α, δ, σ, τ, γ, κ*) is the parameter vector. The RK4 approximation for step *n* is given by

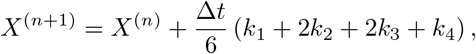

with

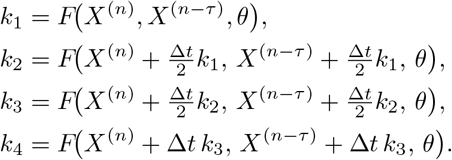

The delay term *X*^(*n*−*τ*)^ was obtained by linear interpolation from previously stored values. The method preserves non-negativity and boundedness of solutions within numerical error.

The forward model was embedded in an outer optimization loop for optimisation routines. Repeated integration under parameter samples drawn from the prior distribution *π*(*θ*) computed the expected utilities. Gradient-free solvers (genetic algorithms and Nelder–Mead simplex) were employed due to the non-convexity of the utility surface. Convergence was assessed by monitoring the stabilization of the objective function value across iterations.

This resolution scheme guarantees numerical stability under the chosen Δ*t* for the parameter ranges in Table 1, as verified by a Courant–Friedrichs–Lewy (CFL) condition check on the fastest dynamics, dominated by *σ* = 0.61.

## 8. Results

### 8.1. Regional Prevalence Dynamics

The metapopulation model captured how cassava brown streak disease spreads over time. Using the prevalence measure, we tracked epidemic trajectories over 180 time units (Figure 2).

**Fig. 1:**
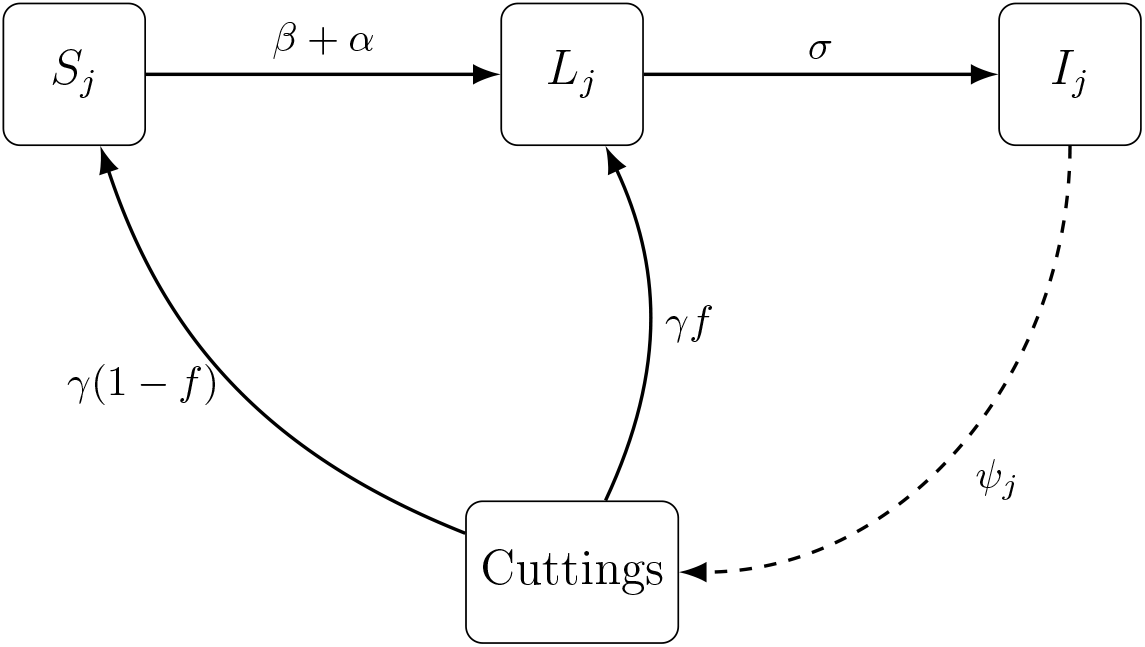
Schematic diagram of the meta-population compartmental model for patch *j*, illustrating transitions between susceptible (*S*_*j*_), latent (*L*_*j*_), and infectious (*I*_*j*_) states in patch *j*compartments. Transmission occurs locally (within the same patch) and non-locally (from other patches) through infection pressure. The diagram also includes recruitment via planting of cuttings, with feedback depending on the infection status of source plants. Delayed progression from latent to infectious states is represented, and feedback loops model the impact of infected cuttings on future susceptibility.

**Fig. 2:**
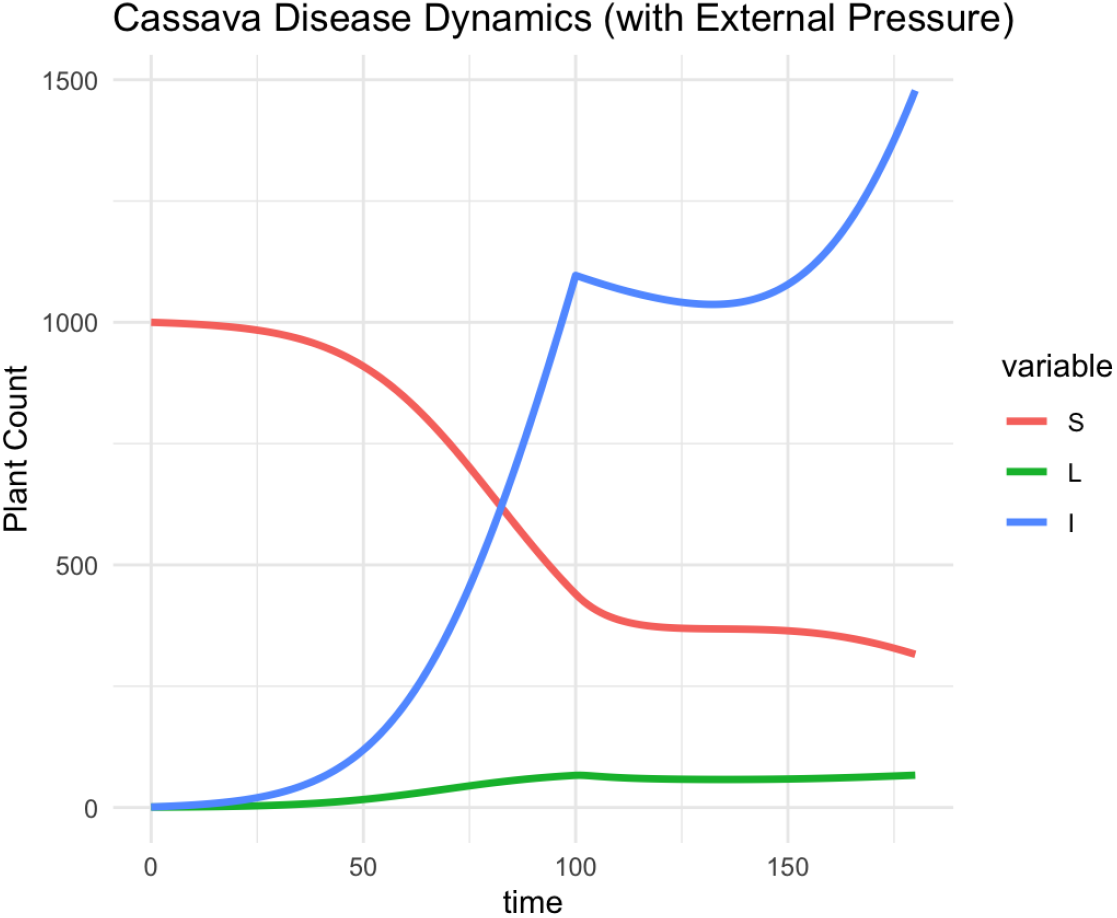
Model-simulated epidemic dynamics of cassava brown streak disease over 180 time units. Susceptible plants decline as infectious plants rise sharply after a delay; latent plants remain low, reflecting the short latent period.

Susceptible plants, starting at 1000, declined steadily as infections accumulated. After a lag of roughly 100 time units, infectious plants surged and eventually outnumbered the susceptible. Latent plants, by contrast, peaked only briefly and remained low throughout, evidencing the model’s short latent period.

This translated into a prevalence curve that initially climbed gradually and then accelerated after the first harvest, when infected seed material re-entered the system. In practice, this reinforces the idea that secondary spread from planting material is a critical driver of epidemic intensification and should be a focal point for intervention and surveillance.

### 8.2. Optimal Sample Size under Different Prevalence Levels

Figure 3 shows that the number of samples depends strongly on prevalence. With high prevalence, only 13 samples were required to achieve a 95% detection probability. At moderate prevalence, this number rose to 31; at low prevalence, it jumped to 50 in the context of parameters (Tables 1).

**Fig. 3:**
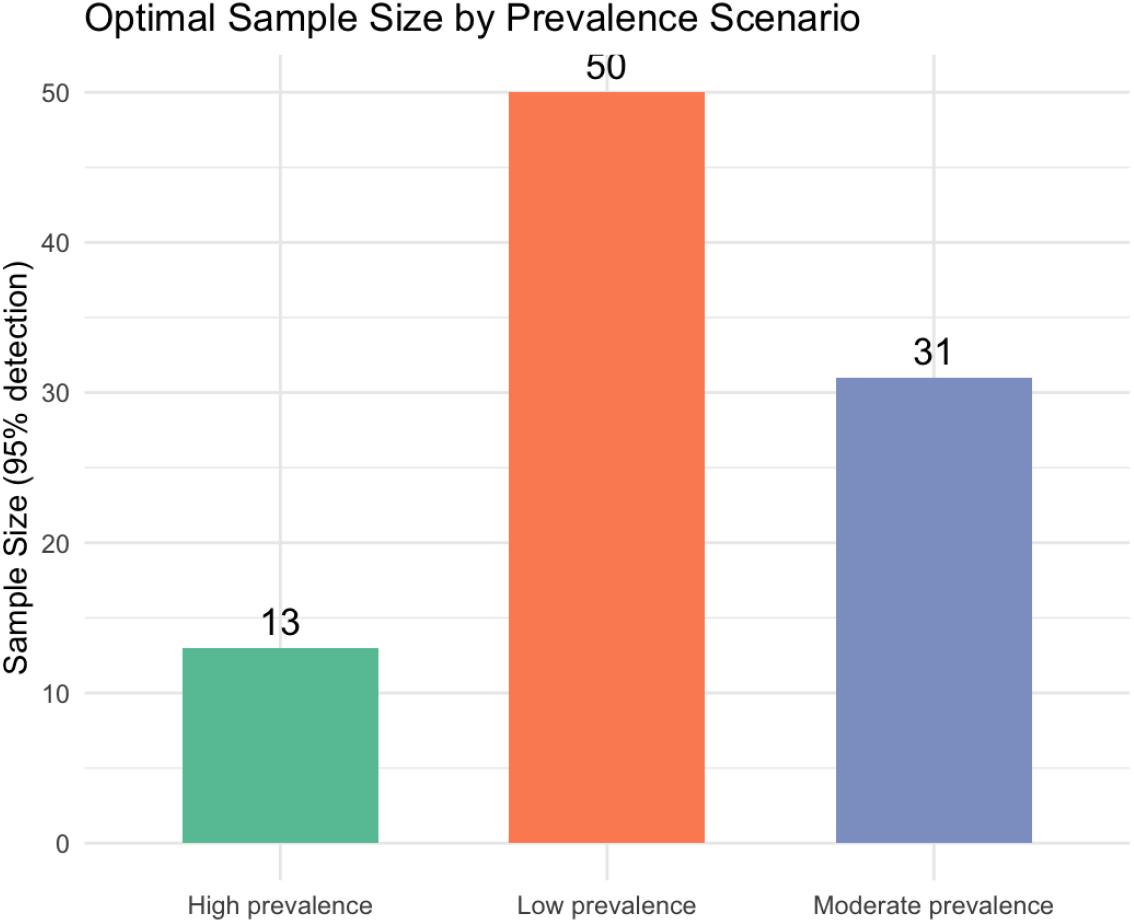
Surveillance effort depends on epidemic stage. The number of samples required for 95% detection increases sharply as prevalence declines, with early detection demanding far greater sampling intensity.

These results underscore a practical challenge: detection is easiest when disease is widespread, but that is also when it is too late. Surveillance becomes far more demanding regarding sample size when prevalence is low, precisely when intervention would be most effective.

### 8.3. Continuous Relationship Between Prevalence and Sample Size

To move beyond three fixed scenarios, we quantified the continuous relationship between prevalence and required sample size (Figure 4). The relationship is distinctly nonlinear. When the prevalence was below 0.1, more than 50 samples were needed. However, once prevalence crossed 0.5, the required effort dropped below 10.

**Fig. 4:**
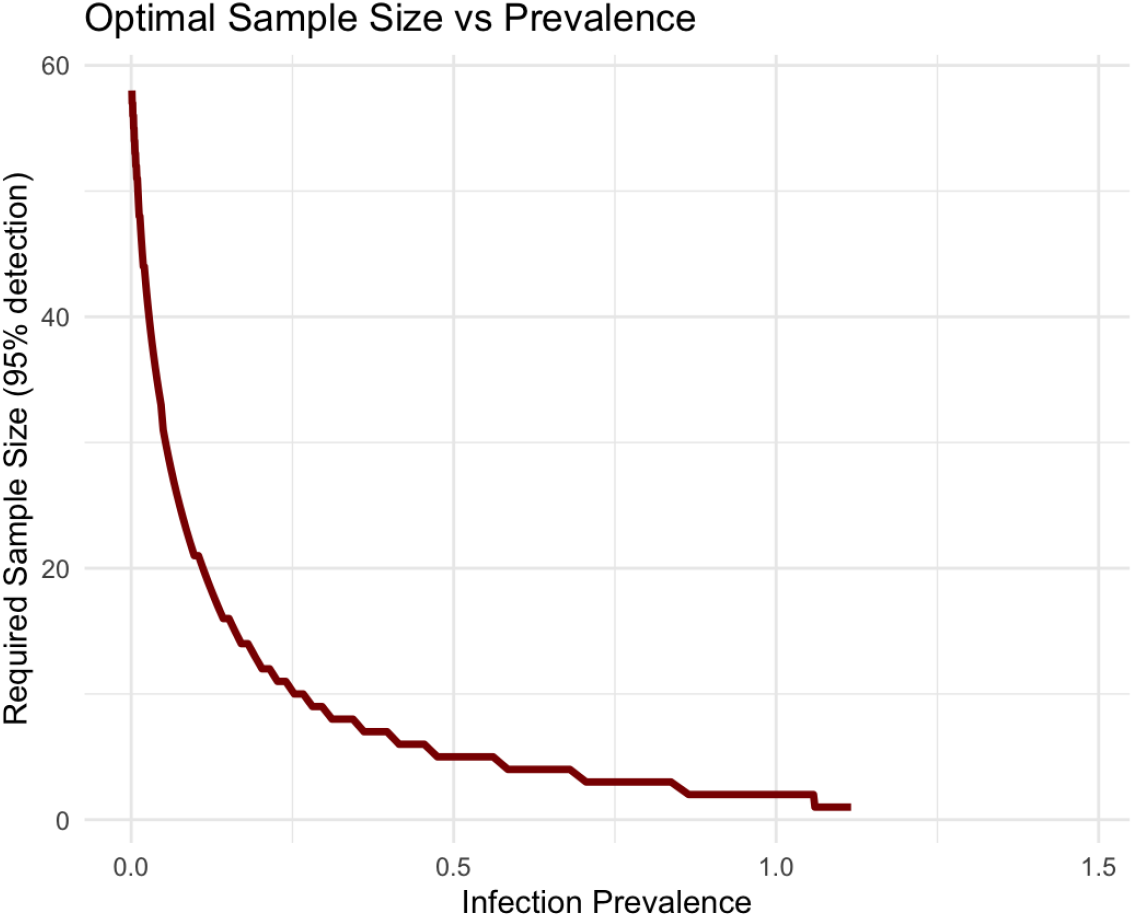
The cost of rarity: sample size requirements decline nonlinearly with increasing prevalence. Detecting early outbreaks (low prevalence) demands disproportionately more effort than established epidemics.

This sharp contrast reveals the cost of rarity: a few percentage points’ difference in prevalence at the low end can double or triple the surveillance burden. For managers, this means that surveillance strategies must budget disproportionately more effort if the goal is to catch outbreaks early.

### 8.4. Sensitivity of Prevalence to Model Parameters

To identify which processes shape epidemic outcomes most strongly, we performed a PRCC sensitivity analysis (Figure 5). Transmission rate (*β*) and progression rate (*σ*) emerged as dominant drivers, both exerting strong positive effects on prevalence. Recovery (*γ*) acted as a counterweight, reducing prevalence. Other parameters contributed modestly, with δ effectively negligible.

**Fig. 5:**
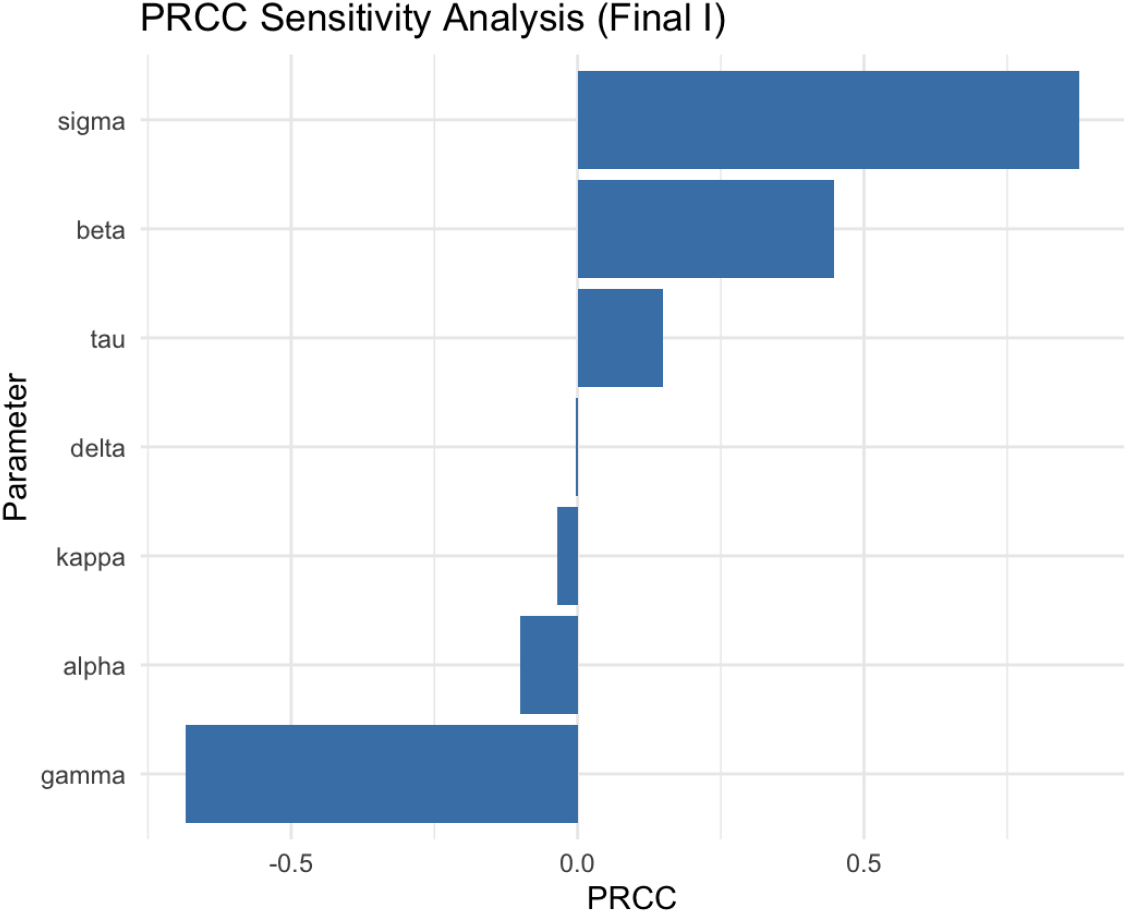
Sensitivity of equilibrium prevalence to model parameters. Transmission (*β*) and progression (*σ*) drive prevalence upward, while recovery (*γ*) reduces it. Other parameters show limited influence.

This ranking suggests that the epidemic hinges on how fast infections are transmitted and how quickly plants progress from latent to infectious. In practical terms, interventions that slow transmission or delay progression will be most effective at bending prevalence curves.

### 8.5. Scenario-Based Spatial Sampling

Finally, we explored how adaptive sequential sampling plays out in space, simulating two contrasting scenarios with 40 farms distributed across Benin. Farm prevalence values of cassava streak disease were assigned randomly between 1-20%, providing a heterogeneous but plausible starting point for the simulations. Spatial interpolation of these baseline values was done using inverse distance weighting (IDW) with a low power parameter (*idp* = 0.1). This setting yields only gentle gradients in the interpolated surface, so adaptive Bayesian updating and sequential decision rules, rather than strong imposed spatial structure, determine the direction of sampling effort (Figure 6).

**Fig. 6:**
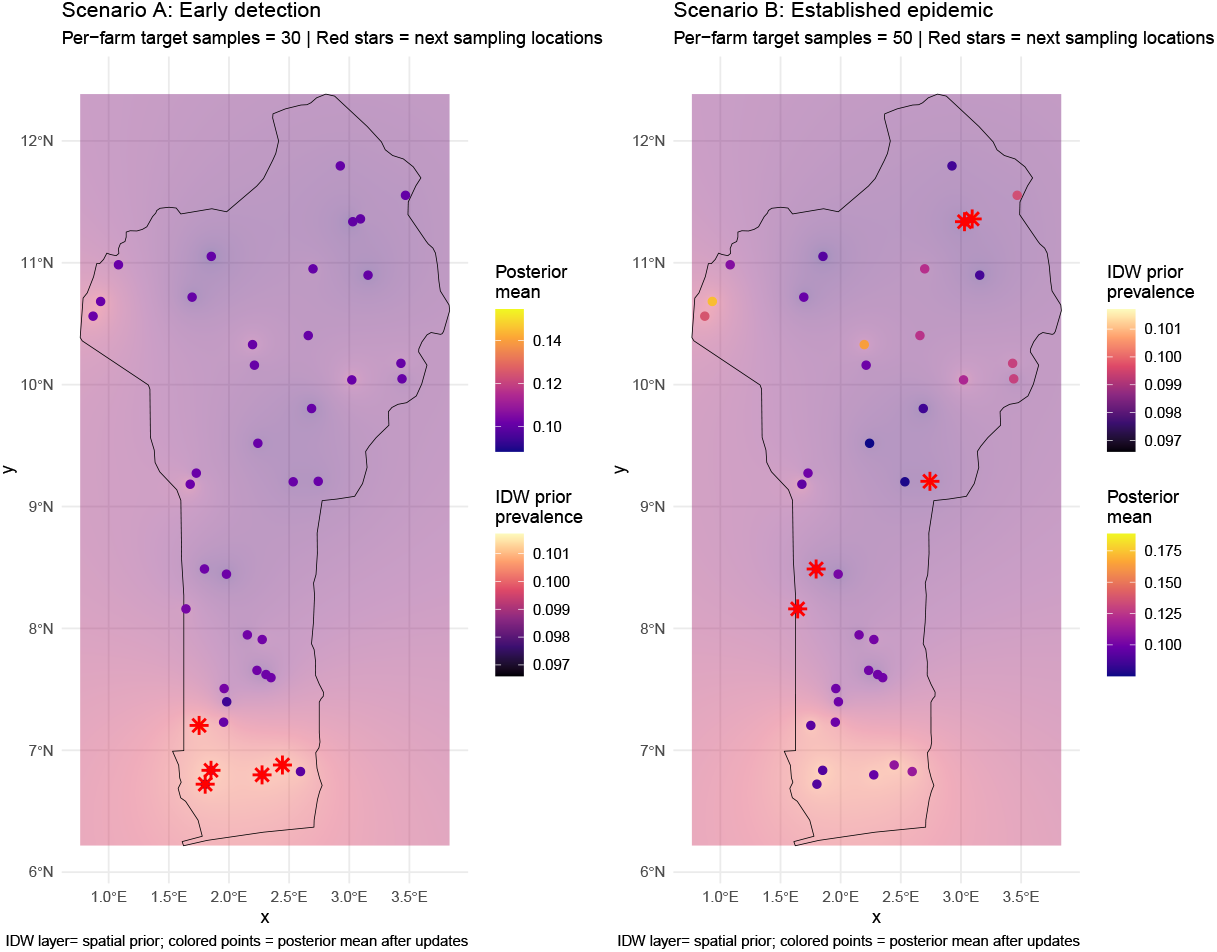
Scenario-based spatial sampling. Farm-level posterior mean prevalence estimates (colored circles) and adaptive resampling priorities (red stars) are shown. A nearly uniform IDW background emphasizes Bayesian updating rather than spatial gradients.

The sequential sampling routine was then applied under two contrasting regimes. In the early detection scenario, farms were allocated up to 30 samples each, and the utility function prioritized sites most likely to exceed the detection threshold (*π*_min_ = 0.05). In the established epidemic scenario, farms were allocated up to 50 samples each, and the sampling utility prioritized the reduction of posterior variance (monitoring). At each round, selected farms received sample batches, posterior distributions were updated, and information was smoothed across the farm network using a power-law dispersal kernel (*η* = 1).

The resulting spatial displays illustrate how effort becomes concentrated adaptively: red stars mark the farms selected for the next sampling round, while the background IDW layer provides the spatial prior (Figure 6). The focus of these maps is not on the interpolation surface itself, but on how Bayesian sequential updates redistribute attention towards farms that contribute most to early detection or reduce uncertainty under established spread.

In Scenario A (early detection, 30 samples per farm) (Figure 6), posterior mean estimates remained close to the prior (centered between 0.05–0.15), but uncertainty was high. Posterior variances were typically on the order of 10^−3^, with some farms showing values nearly twice as large as others. These high-variance farms became the focal points for adaptive sampling, as the algorithm steered new samples toward the most uncertain locations. The IDW surface, though nearly flat, showed a faint yellowish bias in the southern farms, which was reflected in slightly higher posterior means in that region even before strong updates occurred.

In Scenario B (established epidemic, 50 samples per farm) (Figure 6), increased sampling reduced uncertainty dramatically. Posterior variances dropped by more than half compared to Scenario A (often below 510^−4^), and posterior means stabilized closer to the actual prevalence values (0.10–0.18). Adaptive resampling continued to direct effort toward residual high-variance farms, but the landscape picture was already clearer: gradients in prevalence were now apparent, with southern farms tending toward higher posterior means in line with the subtle IDW prior signal.

Unlike the earlier section on optimal sample size, the spatial sampling framework does not determine the required samples. Instead, once sample sizes are fixed, it identifies where those samples are best allocated across the landscape, guided by the IDW prior and refined by Bayesian updating.

Taken together, these scenarios (Figure 6) demonstrate how adaptive sequential sampling changes with epidemic stage. In the early phase, it acts like a spotlight, focusing effort on farms with high variance and uncertain prevalence. Later, with more samples, it works like a fine brush, reducing uncertainty across the board and sharpening gradients, especially where the prior surface, however faint, hints at regional differences such as the slightly elevated prevalence in the south.

### 8.6. Bayesian Estimation

In Scenario A (early detection) (Table 2), farms without sampling defaulted to prior-driven posterior means near 0.10, with high variance (∼ 0.01). Once samples were collected (e.g., farms 20, 24, 34, 37), posterior means adjusted toward observed prevalence and variances shrank sharply (often *<* 0.004), illustrating the power of targeted sampling to reduce uncertainty.

**Table 2:** Comparison of posterior estimates under Scenario A (early detection, 30 samples/farm) and Scenario B (established epidemic, 50 samples/farm). Selected farms illustrate variation in prevalence, sampling intensity, and posterior uncertainty.

| Farm | True prev | Samples | Positives | Posterior mean | Posterior var | Scenario |
| --- | --- | --- | --- | --- | --- | --- |
| 1 | 0.078 | 0 | 0 | 0.101 | 0.0101 | A |
| 20 | 0.141 | 20 | 2 | 0.113 | 0.0035 | A |
| 24 | 0.166 | 30 | 5 | 0.102 | 0.0023 | A |
| 27 | 0.136 | 25 | 1 | 0.088 | 0.0024 | A |
| 34 | 0.165 | 30 | 5 | 0.128 | 0.0029 | A |
| 37 | 0.187 | 30 | 9 | 0.154 | 0.0033 | A |
| 1 | 0.078 | 5 | 0 | 0.099 | 0.0064 | B |
| 9 | 0.164 | 10 | 4 | 0.163 | 0.0072 | B |
| 10 | 0.152 | 5 | 2 | 0.189 | 0.0109 | B |
| 22 | 0.164 | 10 | 5 | 0.172 | 0.0075 | B |
| 34 | 0.165 | 10 | 2 | 0.109 | 0.0051 | B |
| 35 | 0.068 | 5 | 2 | 0.187 | 0.0109 | B |

In Scenario B (established epidemic) (Table 2), smaller per-farm sample sizes (5–10) produced more variable posterior means. Farms with positive detections (e.g., 9, 10, 22, 35) showed posterior means that overshot or undershot the truth depending on the small numbers observed, with variances typically between 0.005 and 0.011. This reflects the efficiency and volatility of inference when many farms are lightly sampled.

Overall, Scenario A demonstrates how concentrated sampling quickly refines estimates at a few targeted farms, while Scenario B spreads effort more thinly, producing a broader but noisier estimate. Bayesian updating remains responsive when positives appear; posterior means shift toward higher prevalence. When negatives dominate, estimates are pulled downward. Variance is the compass for adaptive resampling, highlighting where uncertainty remains greatest.

## 9. Discussion

Modeling disease dynamics of vegetatively propagated species provides a perspective incorporating the complexities of latency and feedback mechanisms within the system. As we illustrate concerning disease prevalence, the initial wave of epidemics tends to be steady and progresses slowly. However, following a latency period, often coinciding with the subsequent planting season characterized by infected planting material, there is a marked increase in prevalence. The increased prevalence can be attributed to the heightened reinfections, as depicted in Figure 2. Our model corroborates the notion that once planting material becomes infected, it may remain asymptomatic during the initial growing season, and in the subsequent planting seasons unfold, the pathogen load within the plants escalates, leading to symptoms, seed degeneration, and ultimately yield loss [4, 20].

Specifically, our analytical framework demonstrated a nonlinear relationship between prevalence and sample size. On the onset of the Epidemic, when prevalence is low, the sampling effort required is disproportionately higher than in areas with moderate to high prevalence. Considering the high level of uncertainty in the detection of infected farms, the investment in time, equipment, and human resources required is higher to decipher the transmission of the Epidemic and design strategic response planning. Nevertheless, the uncertainty and cost associated with passive surveillance via in-field farm surveys could be reduced by leveraging technological advancements and data analytics such as thermography, chlorophyll fluorescence, and hyperspectral sensors combined with a few samples, especially around potential entry routes [14]. These techniques were used elsewhere for early detection of infected individuals even when symptoms are still hidden; they are based on subtle modifications in the plant’s internal temperature, abnormality in photosynthesis, and a change in the range of spectral resolution of leaves [12, 11, 14].

Our research integrates disease prevalence directly into utility-focused frameworks designed to optimize sampling strategies. This approach shows promise for improving decision-making by refining sampling protocols at the initiation and monitoring stages of epidemics in vegetatively propagated crops. Operationally, the framework employs Bayesian updating coupled with spatial interpolation techniques, thereby merging model-based inference with practical decision-making processes. This synergy increases the precision of prevalence estimations and optimizes the allocation of resources and strategic planning. Specifically, the application of Inverse Distance Weighting interpolation has highlighted both considerable benefits and inherent limitations. While IDW improves the clarity of data visualizations, facilitating a better understanding of spatial prevalence patterns and trends for stakeholders, its susceptibility to parameter selection can lead to variability. This sensitivity emphasizes the need for potential enhancements to the geostatistical methodologies employed within the proposed framework.

One of the study’s limitations is failing to account for the erratic pattern of disease often underlying micro and macro environmental conditions while using this framework. As such, our mathematical model describes homogeneous mixing, and we did not account for the horizontal transmission by potent vectors. This oversimplification can lead to an increase in uncertainty in predicting the disease trajectory. Moreover, this analysis did not incorporate landscape-scale dispersal processes, leading to the real-world field distribution. Nonetheless, the framework allows for demonstrating that at the onset of an epidemic, the sampling budget is higher than when the Epidemic is established. It showcases spatial variability in the sampling and where it needs to be surveyed when a data-driven IDW layer is established.

## Acknowledgments

The authors thank the WAVE team, especially Professor Tiendrébéogo Fidèle, for carefully proofreading the manuscript.

